# Network-Based Discovery of Opioid Use Vulnerability in Rats Using the Bayesian Stochastic Block Model

**DOI:** 10.1101/2021.06.30.450568

**Authors:** Carter Allen, Brittany N. Kuhn, Nazzareno Cannella, Ayteria D. Crow, Analyse T. Roberts, Veronica Lunerti, Massimo Ubaldi, Gary Hardiman, Leah C. Solberg Woods, Roberto Ciccocioppo, Peter W. Kalivas, Dongjun Chung

**Affiliations:** Department of Biomedical Informatics, The Ohio State University, Columbus, OH, United States; Department of Neuroscience, Medical University of South Carolina, Charleston, SC, United States; School of Pharmacy, University of Camerino, Camerino, Italy; School of Biological Sciences, Queen’s University Belfast, Belfast, Northern Ireland, United Kingdom; Department of Internal Medicine, Wake Forest University School of Medicine, Winston-Salem, North Carolina, United States

**Keywords:** clustering, community detection, Bayesian modeling, network, opioid use disorder, animal study, individual variation

## Abstract

Opioid use disorder is a psychological condition that affects over 200,000 people per year in the U.S., causing the Centers for Disease Control and Prevention to label the crisis as a rapidly spreading public health epidemic. It has been found that the behavioral relationship between opioid exposure and development of opioid use disorder varies greatly between individuals, implying existence of sup-populations with varying degrees of opioid vulnerability. In this study, we assessed several behavioral variables across heroin taking, refraining and seeking to establish how these factors interact with one another resulting in a heroin dependent, resilient, or vulnerable behavioral phenotype. Over 400 (male and female) heterogeneous stock rats were used in these two studies, and data were collected from two geographically distinct locations. Rats underwent heroin self-administration training, followed by a progressive ratio and heroin-primed reinstatement test. Next, rats underwent extinction training and a cue-induced reinstatement test. To assess how these variables contribute to heroin addiction vulnerability, we developed a network-based data analysis workflow. Specifically, we integrated different cohorts of rats, remove possible batch effects, and constructed a rat-rat similarity network based on their behavioral patterns. We then implemented community detection on this similarity network using a Bayesian degree-corrected stochastic block model to uncover sub-populations of rats with differing levels of opioid vulnerability. We discovered three distinct behavioral sub-populations, each with significantly different behavioral outcomes that allowed for unique characterization of each cluster in terms of vulnerability to opioid use and seeking. We implement this analysis workflow as an open source `R` package, named `mlsbm`.

## 1 Introduction

Opioid addiction is a chronic neuropsychiatric disorder characterized by compulsive drug taking and relapse, despite efforts to remain abstinent. Opioid use disorder (OUD) has risen substantially in the United States over the past two decades, for both prescription drugs (NIH, 2021), as well as illicit opioids, notably heroin (Jones et al., 2015). The parallel rise in both prescription and illicit opioid use and abuse are related to one another, as a majority of heroin users report using prescription opioids prior to heroin use (Cicero et al., 2014; Jones et al., 2015; Compton et al., 2016). Death due to an overdose is also positively correlated between these two opioid classes (Jones et al., 2015), posing an additional obstacle in addressing the current opioid epidemic. Furthermore, heroin use since 2000 has increased in all demographics, regardless of age, sex or socio-economic status (Jones et al., 2015; Compton et al., 2016), suggesting factors independent of these are contributing to the escalation in OUD. This ubiquitous increase in heroin use and dependence across disparate populations highlights the need to assess how individual variation in multiple behavioral traits may be interacting to contribute to an OUD resilient versus vulnerable phenotype.

OUD remains such a critical social and personal problem in part because we do not have animal models that readily predict neurological pathologies for OUD. This problem is compounded by most animal models focusing on one or two behavioral phenotypes, then applying the power of animal experimentation to uncover circuitry and cellular mechanisms for individual phenotypes. In contrast, OUD is a disorder containing many behavioral traits that may contribute differentially to resilience and vulnerability to drug addiction depending on individual genetics and sociology (AP, 2013; Shmulewitz et al., 2015; Venniro et al., 2020). While some studies have created addiction scores that consist of multiple traits, these traits are summed in a linear manner that ignores both individual genetics and environmental exposure (Deroche-Gamonet et al., 2004; Venniro et al., 2018). To overcome these barriers we explore a multidimensional data clustering strategy of 7 behavioral traits characteristic of heroin use and seeking in 451 outbred rats, examined in two distinct laboratories, one at the Medical University of South Carolina in the USA and the other at the University of Camerino in Italy. We sought to uncover differential resilience and vulnerability subgroups within the animal cohort using data clustering methods.

Various types of clustering algorithms have been proposed including k-means clustering (Forgey, 1965), hierarchical clustering (McQuitty, 1966), and mixture modeling (McLachlan et al., 2019). However, behavioral studies generate complex multivariate measurements which can make clustering difficult using standard algorithms. Recently, network-based clustering approaches have become popular across multiple disciplines due to their flexibility and applicability to high-dimensional data. For example, in high dimensional single cell genomics studies, these algorithms are employed in multiple software packages for identifying latent cell types such as T and B cells (Hao et al., 2020). In general, these network-based clustering approaches first construct a similarity network based on observations and then implement a community detection algorithm on this similarity network to identify underlying clusters. As a result, these approaches are less affected by violations of underlying assumptions, such as Gaussianity.

In this paper, we focus on the stochastic block model (SBM), which has strong and rigorous theoretical foundation in statistics literature (Holland et al., 1983; Snijders and Nowicki, 1997). In essence, the SBM allows for identification of latent communities using a probabilistic model describing interconnectivity between nodes within each cluster and between clusters. Due to its probabilistic nature, the SBM has multiple strengths over deterministic algorithmic approaches. First, it provides uncertainty measures for identified clusters, which are critical information to understand the latent structure, e.g., understanding gradual changes across multiple latent clusters (Section 2.6). Second, using goodness-of-fit measures, SBM helps selection of the number of clusters,

which is a long-standing problem in clustering and not straightforward to address in deterministic algorithmic approaches (Section 2.5.2). In addition, the SBM fits naturally into the Bayesian framework, allowing for incorporation of prior expert knowledge to guide the clustering (Snijders and Nowicki, 1997).

This paper is constructed as follows. In Section 2, we develop a Bayesian SBM for animal phenotyping, discuss relevant computational approaches, issues, and software implementation. In Section 3, we apply the proposed methodology to our novel OUD behavioral data. In Section 4, we summarize findings and relevant limitations and future works.

## 2 Methods

### 2.1 Experimental Methods

All experimental procedures were approved by the Institutional Animal Care and Use Committee at the Medical University of South Carolina and by the Italian Ministry of Health (approval 1D580.18). Procedures abided by the National Institute of Health Guide for the Care and Use of Laboratory Animals and the Assessment and Accreditation of Laboratory Animals Care, as well as the European Community Council Directive for Care and Use of Laboratory Animals.

A total of 451 heterogeneous stock (HS: originally n/NIH-HS) rats (male, n=238; female, n=213) bred at Wake Forest University (currently NMcwiWFsm:HS; Rat Genome Database number 13673907) were used in these studies. HS rats were outbred from eight inbred strains and maintained in a way to minimize inbreeding (Hansen and Spuhler, 1984), allowing genetic finemapping to relatively small intervals (Woods and Palmer, 2019). Animals were shipped in batches of 40 (20 males and 20 females per site) to either the Medical University of South Carolina (USA) or the University of Camerino (Italy) at approximately 5 weeks of age. Upon arrival, animals were pair-housed and left undisturbed in a climate-controlled colony room with a standard 12-hour light:dark cycle for 3 weeks prior to the start of testing. Throughout training, rats had *ad libitum* access to food and water. Testing occurred during the dark cycle, between 18:00 h and 6:00 h. Heroin hydrochloride supplied by the National Institute on Drug Abuse (Bethesda, MD) dissolved in 0.9% sterile saline was used in these studies.

Following the 3-week acclimation period, rats underwent surgery under isoflurane anesthesia for the implantation of an indwelling jugular catheter. An analgesic (Ketorolac, 2 mg/kg, sc; or Meloxicam, 0.5 mg/rat, sc), and antibiotic (Cefazolin, 0.2 mg/kg, sc; or enrofloxacin, 1 mg/kg, iv), were administered pre-operatively. Rats were given a minimum of three days of recovery prior to heroin self-administration training commencing. All testing occurred in standard behavioral testing chambers (MED Associates, St. Albans, VT, USA). Presses on an active lever resulted in presentation of a light and tone cue for 5-seconds and an infusion of heroin (20 µg/kg/100 ul infusion) on a fixed-ratio 1 schedule of reinforcement. At the start of the infusion, the house light also turned off for 20-seconds signaling a time-out period during which additional presses on the active lever were recorded but without consequence. Presses on the inactive lever were recorded but without consequence. Sessions lasted for 12 hr or until 300 infusions were earned. Self-administration occurred Monday-Friday, with one session off per week, for a total of four sessions/week. Following 12 self-administration sessions rats underwent a progressive ratio test whereby the number of presses *p*(*t*) required to receive an infusion increased exponentially after each infusion *t* = 1, …, *T* according to the function *p*(*t*) = 5*e*^0.2*t*^ -5 (Richardson and Roberts, 1996). Rats then had three more days of self-administration training to re-establish baseline heroin-taking behavior prior to tests for reinstatement.

At the conclusion of heroin self-administration training, rats underwent a within-session extinction-prime test that lasted for 6 hours. The first 4 hours were extinction training conditions during which presses on both the active and inactive lever were recorded but without consequence (i.e. active lever presses no longer result in presentation of the light/ tone cues or heroin infusion). With two hours left in the session, rats were administered an injection of heroin (0.25 mg/mg, sc), and continued testing under extinction conditions. Daily extinction training sessions (2 hr) then commenced for 6 consecutive days prior to a test for cue-induced reinstatement. During this test, presses on the active lever resulted in presentation of the light/tone cue and turning off of the house light, but no heroin infusions.

At the conclusion of training, several behavioral measures were selected for clustering analyses to reflect the different facets of drug addiction: drug-taking, refraining and seeking behaviors. Heroin-taking behaviors include total heroin consumption (total µg/kg heroin consumed across the first 12 self-administration training session), escalation of intake (total heroin consumed the first three days of self-administration subtracted from the last three days) and break point achieved during the progressive ratio test. The break point is the total number of active lever presses the rat is willing to perform in order to receive an infusion of heroin. Refraining behavior consisted of active lever presses during the first two hours of the within-session extinction prime test (extinction burst) and the last day of extinction training prior to the test for cue-induced reinstatement (extinction day 6). Heroin-seeking behavior is represented by active lever presses during the heroin-prime and cue-induced reinstatement tests.

### 2.2 Data Pre-processing

#### 2.2.1 Batch Correction for Multi-Site Samples

To analyze the MUSC and UCAM cohorts simultaneously, we first performed a visual inspection of possible batch effects between the two study sites. Specifically, we began by concatenating the raw data matrices from each site into an integrated data matrix, where rows corresponded to individual rats and columns correspond to behavioral measures, as described in Section 2.1. Then, to facilitate visualization, we applied the Uniform Manifold Approximation and Projection (UMAP) (McInnes et al., 2018) algorithm to compute 2-dimensional embeddings for each rat. To correct for the apparent batch effect between study sites, we *z*-score transformed each behavioral measure *within study site*. This allowed for analysis of each behavioral measurement on a standardized scale, and, in effect, regressed out unwanted site-specific effects.

#### 2.2.2 Similarity Network Construction

After integrating the behavioral data from each study site as described in Section 2.2.1, we constructed a rat-rat similarity network as follows. First we defined a single parsimonious subset of relevant behavioral measures from the experiments discussed in Section 2.1 using expert knowledge. Here, the goal was to choose variables that reflected the behavioral propensity of each rat for opioid dependence. Next, we computed Pearson’s correlation between each pair of rats using this single parsimonious variable subset. We then formed a rat-rat similarity network, i.e., a collection of nodes and edges, where nodes in the network represent individual rats and edges represent similarities between rats. We placed an edge from each node to its *R* most similar other nodes based on the rat-rat Pearson’s correlation scores. Here, the number of neighbors *R* is a tuning parameter that controls the density of edges in the similarity network. By default, we adopt the widely used heuristic 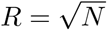 (Stork et al., 2001).

### 2.3 Stochastic Block Model

To detect communities within the overall rat-rat similarity matrix that might correspond to behaviorally distinct sub-populations, we adopted the Bayesian stochastic block model (SBM), a generative model for network data (Snijders and Nowicki, 1997). Let **A** be an *n* × *n* adjacency ma-trix encoding the rat-rat similarity network among *n* total rats, with *A*_*ij*_ = 1 if rat *i* shares an edge with rat *j* (*i* ≠ *j*), and *A*_*ij*_ = 0 otherwise. For a fixed and pre-specified number of communities, *K*, the SBM assumes

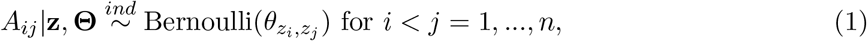

where *z*_*i*_ ∈ {1, …, *K*} is a categorical indicator variable that denotes the community membership of rat *i*, **z** = (*z*_1_, …, *z*_*n*_), and **Θ** is a *K* × *K* connectivity matrix with elements *θ*_*rs*_ described in detail below. Equation (1) implies that the probability of an edge occurring between two nodes depends only on the community membership of each node. Thus, all rats belonging to the same sub-population are regarded as *stochastically equivalent*.

#### 2.3.1 Identifying Community Structures

While our primary object of inference is the vector of latent community indicators **z**, an advantage of the SBM over other community detection algorithms is its ability to conduct statistical inference on the edge probability parameters *θ*_*rs*_, for *r* ≤ *s* = 1, …, *K*. By encoding these parameters in a symmetric connectivity matrix **Θ**, we can obtain a useful characterization of the similarities between communities. Here, diagonal elements of **Θ** are within-community edge probabilities, and offdiagonal elements of **Θ** are between-community edge probabilities. In most cases, we expect to find an *assortative* community structure, in which within-community connections are more likely than between-community connections, though the model is capable of detecting *dissortative* community structures as well (Fortunato and Hric, 2016). Thus, in addition to the community labels, the SBM allows us to characterize the global relationships between communities.

#### 2.3.2 Accommodating Heterogeneous Degree Distributions

Commonly, the SBM as formulated in model (1) is refined to accommodate heterogeneous degree distributions, i.e., *degree correction* (Karrer and Newman, 2011). Since model (1) assumes that the probability of an edge being place between two nodes only depends on the community membership of the nodes, it is not suitable for networks in which each node may have varying degree, that is, the number of edges connected to it. However, as described in Section 2.2.2, our workflow relies on construction of a nearest neighbors network, in which each node, by definition, will have exactly *R* edges, thus degree correction is not necessary.

### 2.4 Prior Specification

We estimate parameters of the SBM using a fully Bayesian approach by assigning prior distributions to all unknown model parameters. We select conjugate priors to obtain closed-form full conditional distributions of all model parameters, which in turn allows for straightforward Gibbs sampling. First, for the cluster indicators *z*_1_, …, *z*_*n*_, we assume a conjugate multinomial-Dirichlet prior with 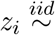 Categorical (π) for *i* = 1, …, *n*, and **π** ∼ Dirichlet(*α*_1_, …, *α*_*K*_), where ***π*** = (*π*_1_, …, *π*_*K*_) control the number of nodes in each community, i.e., the community size. Similarly, we adopt a conjugate beta-Bernoulli prior for **Θ** by letting 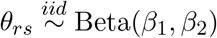 for *r* < *s* = 1, …*K*. By default, we opt for weakly informative priors by setting *α*_1_ = *α*_2_ = … = *α*_*K*_ = 0.5 and *β*_1_ = *β*_2_ = 1 (Gelman et al., 2013).

### 2.5 Posterior Inference

#### 2.5.1 Gibbs Sampler

The model described in Sections 2.3 and 2.4 allows for closed-form full conditional distributions of all model parameters. Thus, we implemented parameter estimation using the following Gibbs sampling algorithm.

1. Update **π** from its full conditional (**π**|**A, z, Θ**) ∼ Dirichlet(*a*_1_, …, *a*_*n*_), where *a*_*k*_ = *α*_*k*_ + *n*_*k*_, and *n*_*k*_ is the current number of nodes in community *k*.
2. For *r* ≤ *s* = 1, …, *K*, update *θ*_*rs*_ from (*θ*_*rs*_ |**A, z, *π***) ∼ Beta(1 + *A*[*rs*], 1 + *n*_*rs*_ - *A*[*rs*]), where *A*[*rs*] are the number of observed edges between communities *r* and *s*, and *n*_*rs*_ = *n*_*r*_ *n*_*s*_ - *n*_*r*_ *I*(*r* = *s*) are the number of possible edges between communities *r* and *s*, and *I*(*r* = *s*) is the indicator function equal to 1 if *r* = *s* and 0 otherwise.
3. For *i* = 1, …, *n*, update *z*_*i*_ from (*z*_*i*_ |*z*_*-i*_, **A**, ***π*, Θ**) ∼ Multinomial(***ρ***_*i*_), where ***ρ***_*i*_ = (*ρ*_*i*1_, …, *ρ*_*iK*_) and 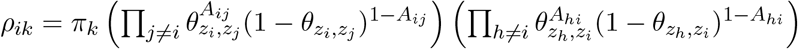.

#### 2.5.2 Choosing the Number of Communities

A critical step of our proposed workflow for identifying behavioral sub-populations in rats is the choice of *K*, i.e., the number of communities. Since the choice of *K* should consider both expert knowledge and evidence from the data, we refrain from proposing a “one size fits all” globally optimal method for choosing of *K*. Instead, in Section 3 we discuss how Bayesian Information Criterion (BIC) (Schwarz et al., 1978) can be used in conjunction with biological knowledge to make informed choices for *K*.

#### 2.5.3 Label Switching

Label switching is an issue encountered in Markov chain Monte Carlo (MCMC) methods, such as the Gibbs sampler proposed above, wherein the model likelihood is invariant to permutations of a latent categorical variable such as **z**. As a result, we may observe natural permutations of **z** over the course of the MCMC sampling that cause the estimates of all other community-specific parameters to be conflated, thereby jeopardizing the accuracy of model parameter estimates. This problem is exacerbated when communities are not well separated. Previous works have attempted to address the issue by re-shuffling posterior samples after the sampling has completed (Papastamoulis, 2016). However, these post-sampling methods rely on prediction and thereby are fallible to prediction error. To address label switching, we adopt the canonical projection of **z** proposed by Peng and Carvalho (2016) in the context of Bayesian SBMs, in which we restrict samples of **z** to the canonical sub-space *Ƶ* = {**z** : ord(**z**) = (1, …, *K*)}. In other words, we permute **z** at each MCMC iteration such that community 1 appears first in **z**, community 2 appears second in **z**, *et cetera*. Finally, we choose as our final estimate of **z** the maximum *a posteriori* (MAP) estimate of of **z** across all post-burn MCMC samples (Gelman et al., 2013).

### 2.6 Continuous Phenotyping

While the SBM presented thus far assumes that the overall experimental cohort can be decomposed into a fixed number of discrete communities, where each experimental unit (e.g., rat) is assigned to exactly one community, often interest lies in further differentiating members within a community in a more continuous fashion. Indeed, a core benefit of the Bayesian SBM is that the discrete model structure may be augmented using uncertainty measures, i.e., a quantification of our inferred level of confidence in each estimated model parameter. For instance, let 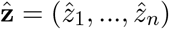 be the posterior estimate of the true SBM community labeling vector **z** obtained from the MCMC estimation procedure described in Section 2.5. Letting *s* = 1, …, *S* index the post burn-in MCMC iterations, we may quantify the uncertainty in each estimate 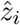 as

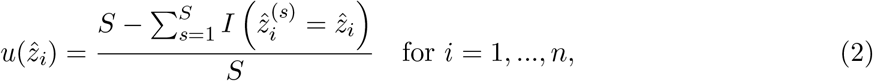

where 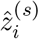 is the estimate of *z*_*i*_ at MCMC iteration *s*, and 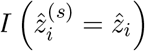 is the indicator function equal to 1 if 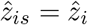 and 0 otherwise. In words, 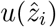 represents the proportion of MCMC iterations where the estimate of *z*_*i*_ was not the posterior MAP estimate 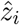. For nodes that share many edges with other nodes within their respective community, i.e., those that are highly typical of their community, the uncertainty measure should be low. Meanwhile, for nodes that share edges with nodes outside of their respective community, the uncertainty measure should be high, as these nodes will likely be assigned to other communities intermittently over the course of the MCMC estimation. In this way we may augment the cluster labels obtained by the SBM with quantification of our level of confidence in them – a significant advantage over other non-model-based clustering methods.

In addition to uncertainty quantification, we may similarly use the MCMC draws 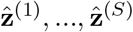 to conduct *continuous phenotyping*, or the ranking of subjects based on their affinity towards a certain phenotype. For example, in our context of assigning rats to vulnerable and resilient phenotypes using the SBM, we may also provide a continuous measure of affinity towards the vulnerable phenotype for each rat that can be used to rank rats within clusters. In this setting, let cluster *k*_*v*_ ∈ {1, 2, …, *K*} be the cluster annotated as vulnerable for opioid dependence. For each rat *i* = 1, …, *n*, we define the continuous phenotype vulnerability score *v*(*i*) as 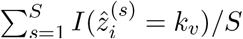, i.e., the proportion of MCMC iterations in which rat *i* is assigned to cluster *k*_*v*_.

### 2.7 Software Implementation

For convenient implementation of the workflow proposed throughout Section 2, we developed `mlsbm`, an efficient and user-friendly `R` package for the identification of sub-populations in network data (Allen and Chung, 2021). The `mlsbm` package is freely available for download from the Comprehensive `R` Archive Network (R Core Team, 2020) at https://cran.r-project.org/package=mlsbm. The `mlsbm` package includes robust documentation to facilitate applications to a variety of clustering tasks.

### 2.8 Comparison to Alternative Approaches

We sought to assess the performance of the SBM clustering workflow relative to alternative clustering approaches, we applied five popular clustering algorithms, namely the Louvain, walktrap, hiearchical clustering, K-means, and DBSCAN algorithms. The Louvain (Blondel et al., 2008) and walktrap (Pons and Latapy, 2005) algorithms, like the SBM, are graph-based methods that operate on the nearest neighbors network described in Section 2.2.2. The Louvain algorithm seeks to maximize the modularity of the graph, a measurement of the strength of clustering structure of a graph relative to randomly generated graphs. The walktrap algorithm uses random walks on the nearest neighbors graph to find the most densely connected sub-graphs, i.e., clusters, within the graph. Hierarchical clustering (McQuitty, 1966) is a “bottom up” approach that iteratively merges the most similar observations into clusters to form a tree structure that can be used to produce cluster labels for a pre-specified value of *K*. K-means (Forgey, 1965) and DBSCAN (Ester et al., 1996) seek to place boundaries around observations in high-dimensional space such that the data points within boundaries, i.e., clusters, are more similar than those across boundaries. While these approaches are commonly used, they lack the inferential benefits of the SBM such as the ability to choose *K* using model fit criteria and provide uncertainty quantification in addition to cluster labels.

## 3 Results

The overall sample was composed of *N*_*m*_ = 238 males and *N*_*f*_ = 213 females. The MUSC study site contributed 243 rats, while the UCAM study site contributed 208. As seen in Figure 1A, the MUSC and UCAM cohorts exhibit clear separation on the 2-dimensional UMAP space, indicating the potential of study site to act as a confounding variable in our analysis, and preventing simultaneous analysis of rats from both cohorts. In Figure 1B, we present the 2-dimension UMAP embedding of the concatenated *z*-score transformed data set, in which no distinguishable separation exists between the MUSC and UCAM rats. Hence, the site-specific z-scoring approach detailed in Section 2.2.1 was able to effectively remove the site-specific batch effect from the data.

**Figure 1:**
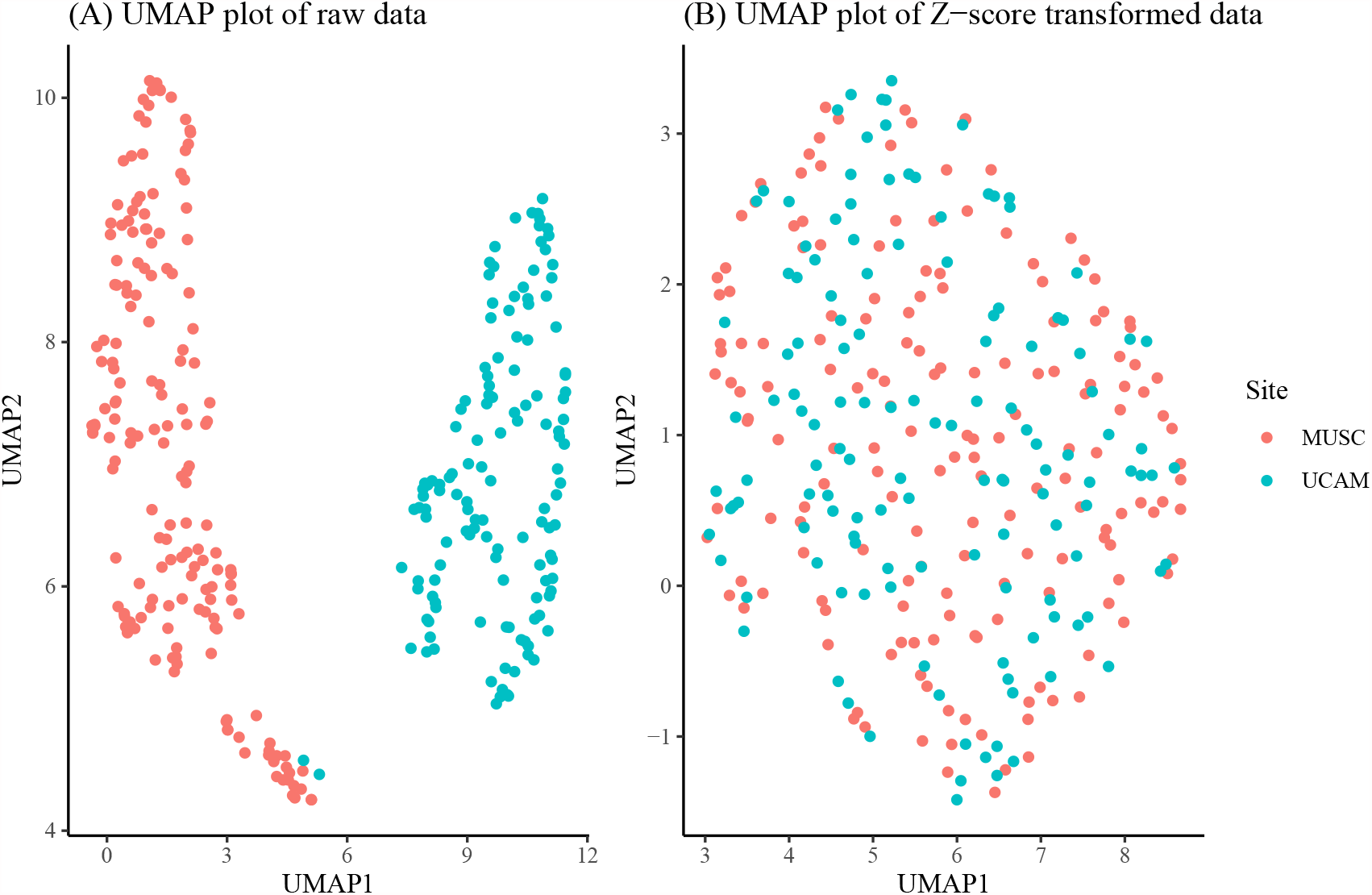
Panel (A): UMAP dimension reduction of behavioral measures before site-specific z-scoring shows significant batch effect of study site (MUSC vs. UCAM). Panel (B): UMAP dimension reduction after site-specific z-scoring shows adjustment for study site batch effect.

To construct the rat-rat similarity network, we computed the correlation between each pair of rats using the 7 variables discussed in Section 2.1 and then formed an adjacency network where each rat was connected to its 21 most similar rats. We applied the SBM clustering analysis described in Section 2.3 to the analysis of *N* = 451 rats. To choose the most appropriate number of clusters *K*, we fit the SBM to the adjacency network for a range of *K* from *K* = 2, …, 10. We ran each model for 10,000 MCMC iterations and discarded the first 1,000 iterations as burn-in, resulting in a total run time of under 4 minutes for each model using a single 4.7 GHz Intel i7 processor. Using BIC, we found that *K* = 3, 4, 5 provided approximately equal goodness of fit, with *K* = 2 or *K >* 5 provided relatively poor fit (Figure 2A). As such, we chose *K* = 3 to provide the most parsimonious representation of the data and to assess the vulnerable, intermediate, and resilient sub-type hypothesis discussed in Section 1. An adjacency matrix with rows and columns sorted by inferred cluster indicators from the 3 cluster model is shown in Figure 2B. Figure 2C shows the SBM estimated cluster labels on UMAP space.

**Figure 2:**
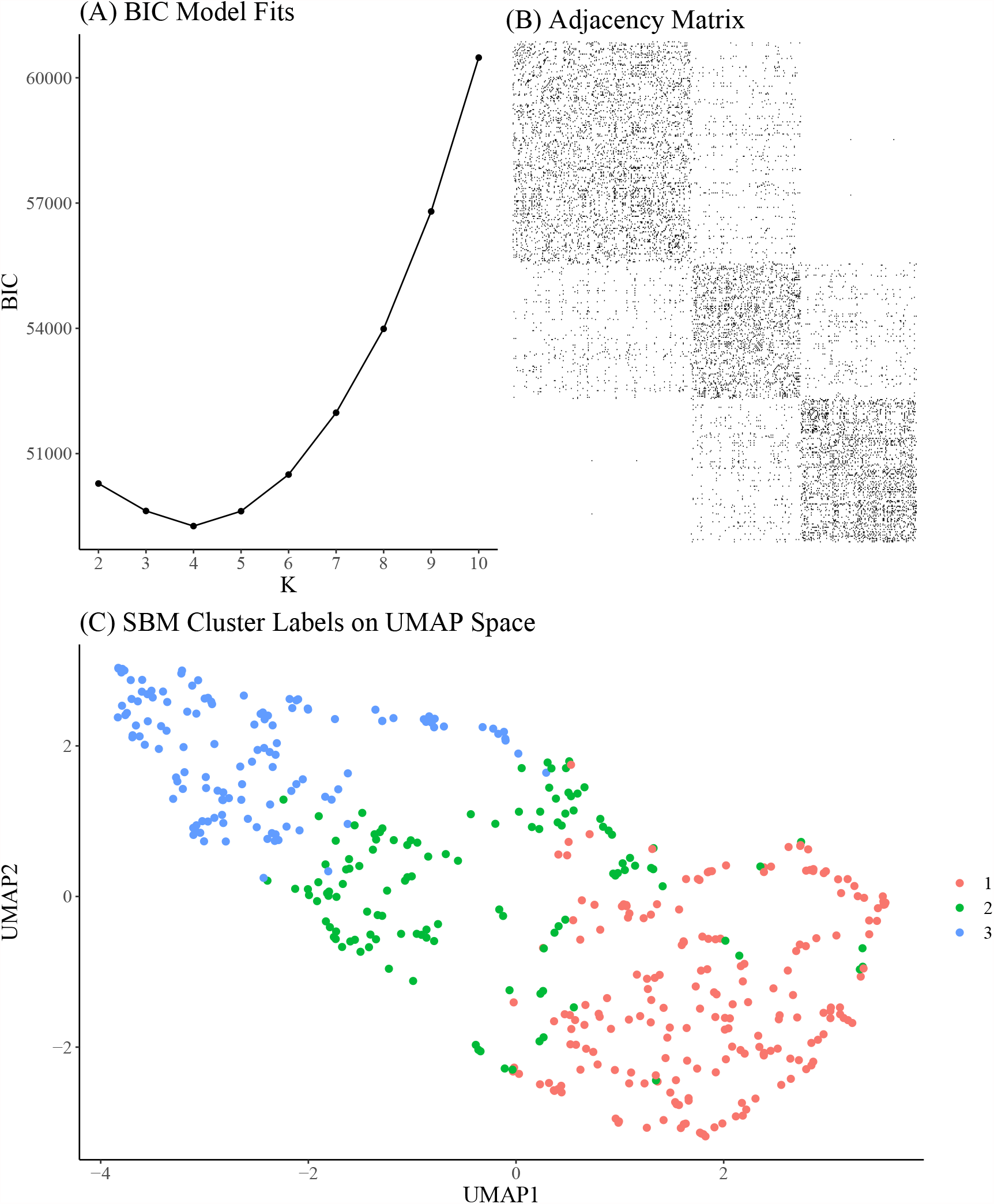
(A) Bayesian Information Criterion (BIC) from SBMs fit with a range of *K. K* = 3, 4, 5 seem to provide similarly optimal fit in terms of BIC. (B) Adjacency matrix of inferred clusters from the SBM using *K* = 3 clusters. (C) UMAP reduction of behavioral measurements colored by inferred cluster labels from the SBM using *K* = 3 clusters.

Figure 3 shows empirical means and 95% *z* confidence intervals for each of the 7 selected behavioral measures across each of the inferred clusters from the SBM. Notably, each cluster appears to show clear separation in most of the behavioral variables. For instance, the total heroin consumption was highest in cluster 1 and lowest in cluster 3, with cluster 2 falling in between clusters 1 and 3, and all 95 % confidence intervals not overlapping. Similarly, cluster 1 demonstrated a more rapid escalation of heroin intake relative to clusters 2 and 3. We quantified the difference between clusters by fitting a one-way ANOVA for each of the 7 behavioral measures vs the SBM cluster indicators. We conducted a global F test for mean differences among groups. F-statistics and associated p-values are displayed in Table 1.

**Figure 3:**
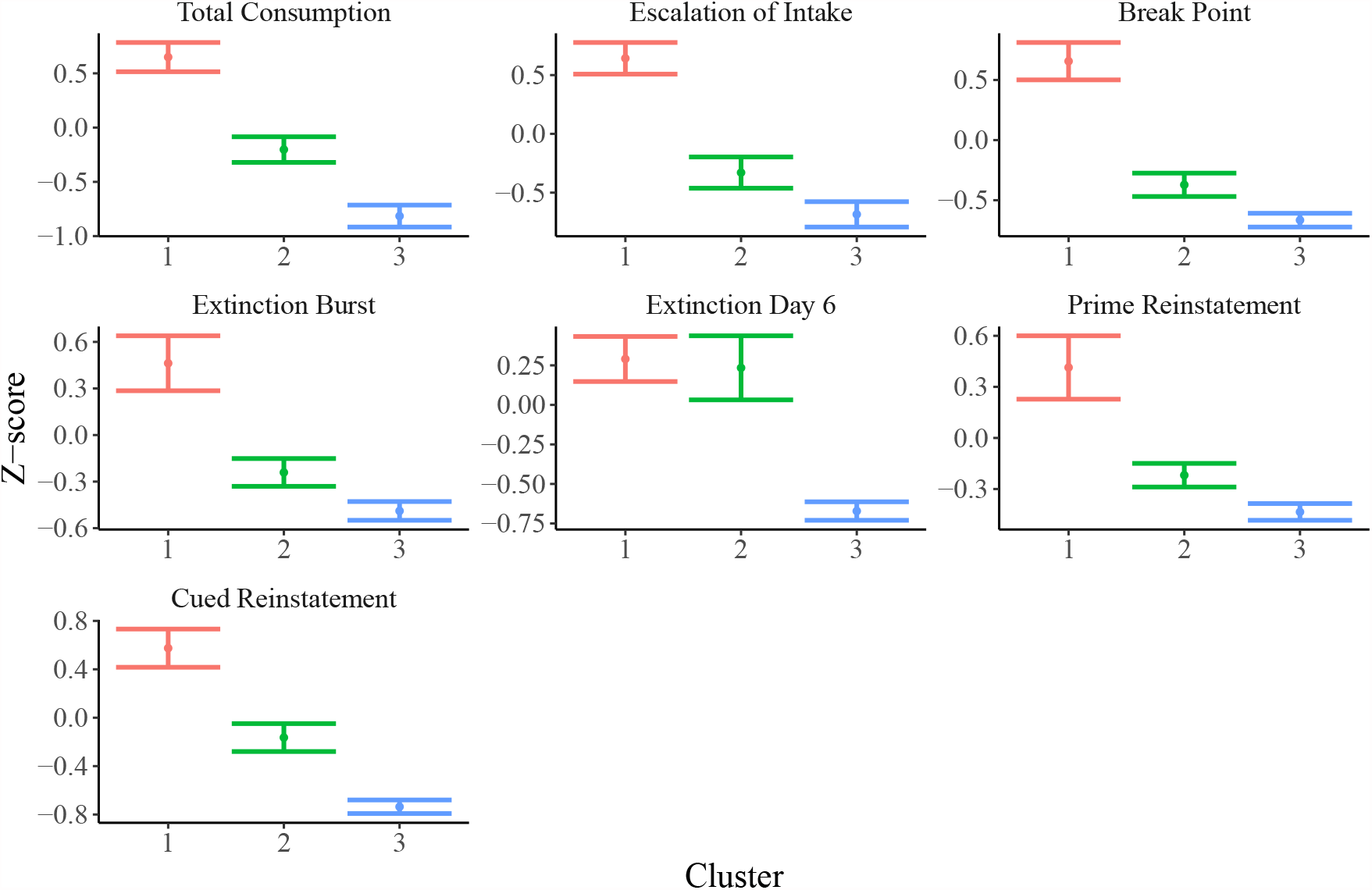
Means and 95% confidence intervals for relevant behavioral measures in each cluster. Distributions of behavioral variables indicate evidence for vulnerable (cluster 1; *N* = 200), inter-mediate (cluster 2; *N* = 122), and resilient (cluster 3; *N* = 129) sub-populations.

**Table 1:** ANOVA global F-statistics and associated p-values for each behavioral measure.

| Variable | F-statistic | P-value |
| --- | --- | --- |
| Total Consumption | 283.8 | < 0.0001 |
| Escalation of Intake | 220.7 | < 0.0001 |
| Break Point | 221.6 | < 0.0001 |
| Extinction Burst | 94.78 | < 0.0001 |
| Extinction Day 6 | 77.12 | < 0.0001 |
| Prime Reinstatement | 72.36 | < 0.0001 |
| Cued Reinstatement | 200.6 | < 0.0001 |

To further investigate the vulnerable, intermediate, and resilient sub-type hypothesis, we leveraged the inferential abilities of the Bayesian SBM to infer the similarity among rats from each cluster. Specifically, by investigating the posterior distribution of the elements of the matrix **Θ**, we may characterize the similarity among rats within and between each of the three clusters. In Figure 4, we show a heatmap of posterior means and 95% Bayesian credible intervals for *θ*_11_, *θ*_22_, *θ*_33_, *θ*_12_, *θ*_13_, and *θ*_23_. We found that the estimated values of the within-cluster connectivity parameters *θ*_11_, *θ*_22_, *θ*_33_ were found to be significantly higher than those of the between-cluster parameters *θ*_12_, *θ*_13_, and *θ*_23_.

**Figure 4:**
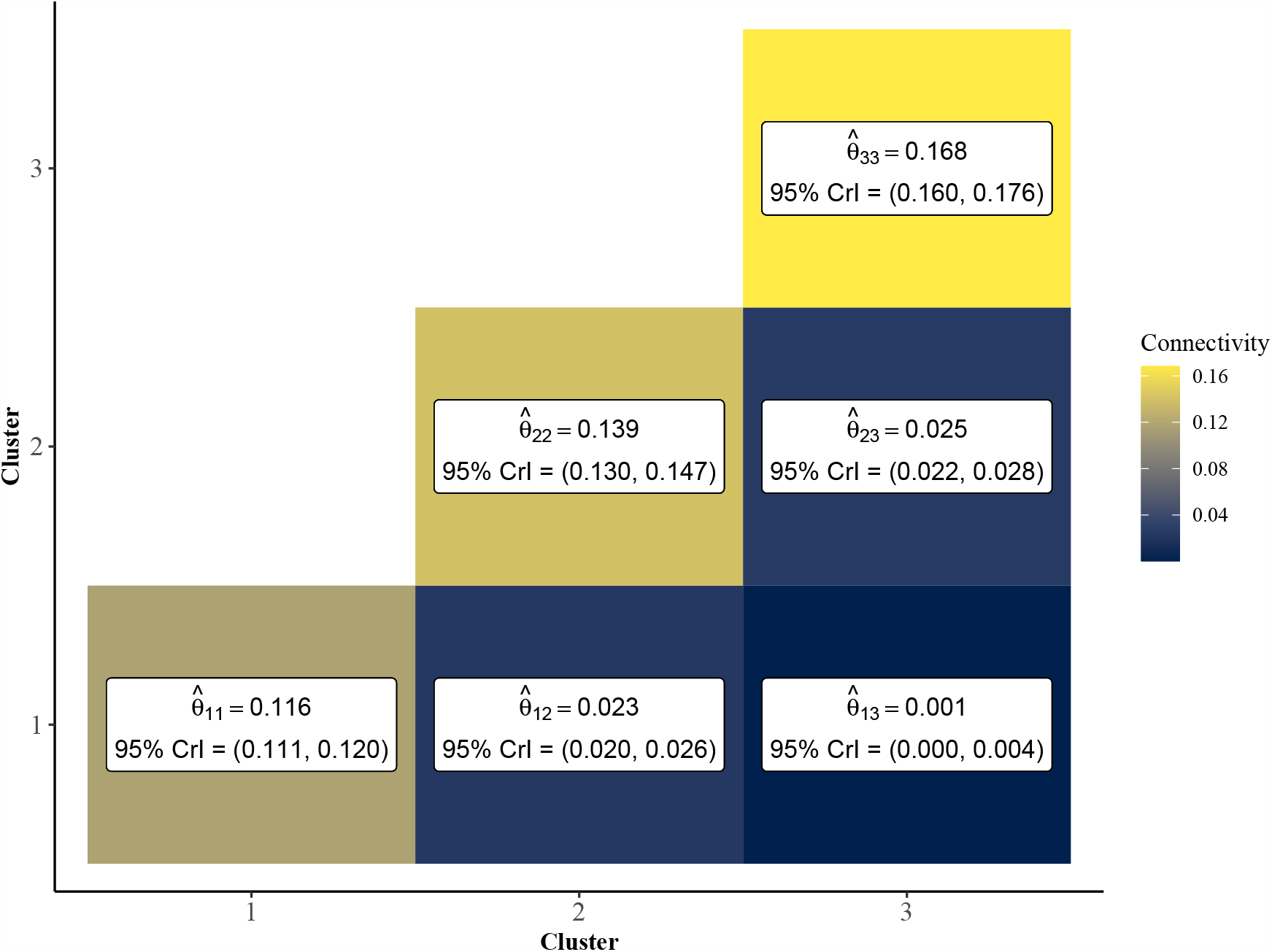
Point estimates and 95% credible intervals of cluster connectivity parameters **Θ**. The SBM estimates higher values for within-cluster connectivity parameters, *θ*_11_, *θ*_22_, and *θ*_33_, which is indicative of an assortative community structure. Thus, rats within the same community are expected to have significantly higher similarity than rats of different clusters. Clusters 1 and 3 are most dissimilar as evidenced by lower values of 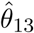 relative to 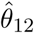 and 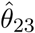.

In fact, cluster 1, which had the weakest estimated within-cluster connectivity 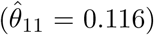, was still over four times more densely connected than the highest between-cluster connection, which was shared between clusters 2 and 3 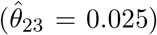. This is indicative of strong assortative community structure in the rat-rat similarity network, in which rats of the same community are more likely to be correlated in terms of behavioral measurements than rats of differing communities. Further, Figure 4 shows that clusters 1 and 3 were the most dissimilar, with cluster 2 serving as an intermediate cluster.

In Figure 5, we plot results from the uncertainty measure and continuous phenotyping analysis presented in Section 2.6. Figure 5A plots the cluster assignments on UMAP space, where each point is sized proportionally to its uncertainty measure of cluster assignment (larger points imply higher uncertainty). We label the ID of each rat that featured an uncertainty measure above 0.10, corresponding to rats that spent at least 10% of the post burn-in MCMC iterations from the *K* = 3 SBM in a cluster other than the cluster it was assigned to by the MAP estimate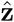. A number of interesting patterns emerge from this uncertainty analysis. First, we find that rats with higher uncertainty tend to be located near borders between clusters on the UMAP space. Interestingly, rat 101, which was assigned to cluster 2 but is surrounded in UMAP space by rats in cluster 3, featured high uncertainty. Meanwhile, several cluster 2 rats were surrounded by cluster 1 rats in the UMAP space but featured low uncertainty.

**Figure 5:**
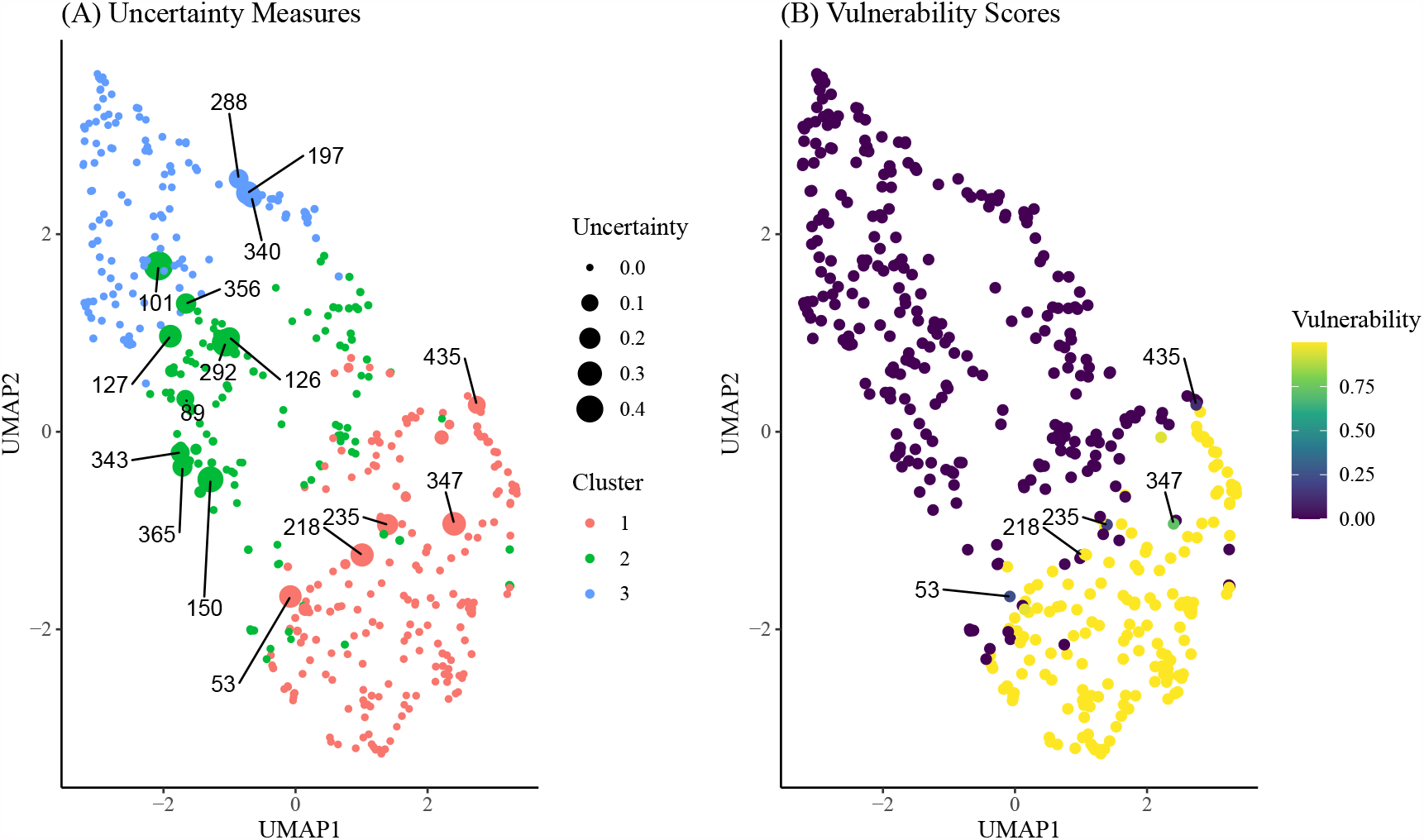
(A) Uncertainty scores of cluster assignment for each rat on UMAP space. Animal IDs are given for subjects with uncertainty measure above 0.10, which is indicative of at least 10% of MCMC iterations spent in a cluster other than the final inferred cluster. (B) Vulnerability scores for each rat on UMAP space. Animal IDs were shown for subjects with uncertainty above 0.10 and vunerability less than 0.90.

Figure 5B displays results from the continuous phenotyping analysis, wherein cluster 1 was annotated as the vulnerable cluster (Figure 3) and chosen as the phenotype of interest. We computed the vulnerability score of each rat as the proportion of post burn-in MCMC iterations from the *K* = 3 SBM that were spent in cluster 1. We labeled the IDs of the most interesting rats: those with uncertainty measures above 0.10 but vulnerability measures less than 0.90. These rats were located on the border between the intermediate cluster 2 and the vulnerable cluster 1, indicating higher propensity towards opioid dependence than other rats in cluster 2. These results demonstrate the ability of continuous phenotyping to augment the clustering results of the SBM to allow for disambiguation of within-cluster differences between subjects.

Figure 6 displays results from alternative clustering methods as described in Section 2.8. The graph clustering Louvain and walktrap algorithms tended to produce a larger number of clusters, each smaller in size relative to the SBM. Due to this, the agreement between the results from these methods and those from the SBM is low (ARI *<* 0.30). Both the hierarchical clustering method using squared Ward dissimilarity (Murtagh and Legendre, 2014) and the K-means algorithm resulted in moderate agreement with the SBM (ARI = 0.343 and 0.374, respectively), while the DBSCAN algorithm yielded a 4 cluster result using default parameters, two of which were sparsely populated. These results suggest the SBM is best suited to addressing the research question at hand.

**Figure 6:**
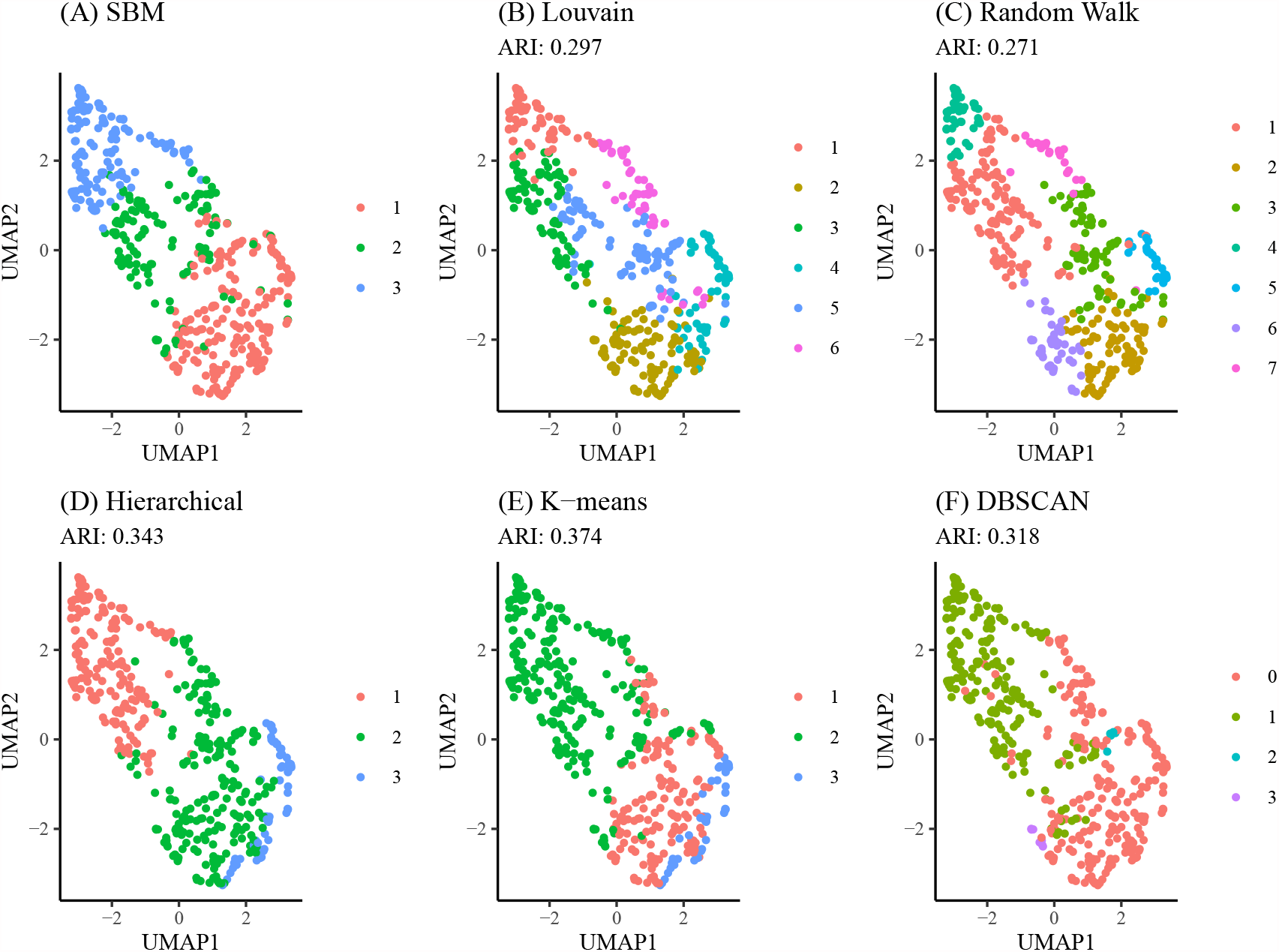
Comparison of SBM performance relative to alternative methods. (A) Clustering results from the SBM using *K* = 3. (B) Clustering results from the Louvain algorithm (no tuning parameters available). (C) Clustering results from the Walktrap algorithm using random walks of length 4. (D) Hierarchical clustering results using a dendrogram cut at *K* = 3. (E) K-means clustering results using the Hartigan-Wong method and *K* = 3. (F) DBSCAN clustering results using a radius of 0.8 and minimum neighborhood size of 5

## 4 Conclusion

We have developed a comprehensive community detection framework for the analysis of behavioral sub-populations in a cohort of 451 outbred rats subject to heroin self-administration exposure. We discovered the presence of batch effects between the two study sites that contributed to this cohort, and we corrected for these effects using study-site specific z-scoring. Seven behavioral measures were chosen to characterize the vulnerability of each rat to forming opioid dependence. Taken together, these measures quantified three important aspects of dependence: drug-taking, refraining and seeking behaviors. Using these measures, we then converted the multidimensional behavioral data into a rat-rat similarity network, which allowed for investigation of distinct communities within the overall network.

We chose the Bayesian stochastic block model, a statistical model for network data, for inves-tigation of behavioral sub-populations within this cohort. We used the model fit criterion BIC to choose a subset of best fitting models in terms of number of communities. Of this best fitting subset, we chose the three cluster model as it offered the best balance between optimizing statistical and biological criteria. Using ANOVA global F-tests, we found significant separation between clusters in terms of each of the seven behavioral measures. Additionally, investigation of average trends across clusters in each behavioral measure allowed us to annotate vulnerable, resilient, and intermediate sub-groups with high confidence. Using the community connectivity parameters inferred by the SBM, we described the relative similarity between clusters, with the vulnerable and resilient clusters each displaying similarity to the intermediate cluster but very little similarity to one another.

To augment the discrete community labels obtained from the SBM, we developed an uncertainty measure, which uses samples from the posterior distribution of the cluster labels to estimate our confidence in the inferred community structure. We also implemented continuous phenotyping to investigate heterogeneities within clusters in terms of vulnerability to opioid dependence. We found a subset of intermediate vulnerability animals who featured relatively high affinity towards the vulnerably cluster, providing candidate animals for further investigation of the differences between vulnerable and resilient animals. Finally, we developed `mlsbm`, an efficient and robust `R` package for implementation of our proposed clustering workflow. The `mlsbm` package is publicly available through CRAN for use in future behavioral studies.

This work can be extended in a number of ways. First, in the presence of explanatory covariates related to but not including behavioral measures, we may implement statistical models such as multinomial regression on cluster indicators to further explain cluster allocation. However, we reserve this for future work as no such covariates were available for our analysis. Second, pending the completion of ongoing experiments, we plan to develop integrative analysis frameworks to jointly model behavioral data with other modalities such as genetic sequencing data to infer correlations between behavioral phenotypes and genotypes of rats. Finally, using our existing `mlsbm` software, we plan to develop an interactive web application to analyze a variety of network-based data sets without the need for programming experience in `R`.

## Author Contributions

The behavioral experiments were designed by NC, BNK, MU, LSW, GH, RC and PWK. All behavioral experimental procedures were conducted by BNK, NC, VL, ADC and ATR. Statistical modeling, software development, and data analyses were conducted by CA and DC. This manuscript was written by CA, BNK, NC, RC, PWK and DC.

## Data Availability Statement

The `R` package `mlsbm` is publicly available from the Comprehensive `R` Archive Network https://cran.r-project.org/package=mlsbm. The behavioral data used in this paper are not readily available due to ongoing data collection, which is implemented as part of the ongoing NIH-funded research project. Please contact the corresponding author for any inquiry related to the behavioral data.

## References

Allen, C. and Chung, D. (2021). mlsbm: Efficient Estimation of Bayesian SBMs & MLSBMs. R package version 0.99.2.

AP, A. (2013). Diagnostic and statistical manual of mental disorders. Am Psychiatric Assoc, Fifth Edition 21,.

Blondel, V. D., Guillaume, J.-L., Lambiotte, R., and Lefebvre, E. (2008). Fast unfolding of communities in large networks. Journal of statistical mechanics: theory and experiment 2008, P10008.

Cicero, T. J., Ellis, M. S., Surratt, H. L., and Kurtz, S. P. (2014). The changing face of heroin use in the united states: a retrospective analysis of the past 50 years. JAMA psychiatry 71, 821–826.

Compton, W. M., Jones, C. M., and Baldwin, G. T. (2016). Relationship between nonmedical prescription-opioid use and heroin use. New England Journal of Medicine 374, 154–163.

Deroche-Gamonet, V., Belin, D., and Piazza, P. V. (2004). Evidence for addiction-like behavior in the rat. Science 305, 1014–1017.

Ester, M., Kriegel, H.-P., Sander, J., Xu, X., et al. (1996). A density-based algorithm for discovering clusters in large spatial databases with noise. In Kdd, volume 96, pages 226–231.

Forgey, E. (1965). Cluster analysis of multivariate data: Efficiency vs. interpretability of classification. Biometrics 21, 768–769.

Fortunato, S. and Hric, D. (2016). Community detection in networks: A user guide. Physics reports659, 1–44.

Gelman, A., Carlin, J. B., Stern, H. S., Dunson, D. B., Vehtari, A., and Rubin, D. B. (2013). Bayesian data analysis. CRC press.

Hansen, C. and Spuhler, K. (1984). Development of the national institutes of health genetically heterogeneous rat stock. Alcoholism: Clinical and Experimental Research 8, 477–479.

Hao, Y., Hao, S., Andersen-Nissen, E., Mauck, W. M., Zheng, S., Butler, A., Lee, M. J., Wilk, A. J., Darby, C., Zagar, M., et al. (2020). Integrated analysis of multimodal single-cell data. bioRxiv.

Holland, P. W., Laskey, K. B., and Leinhardt, S. (1983). Stochastic blockmodels: First steps. Social networks 5, 109–137.

Jones, C. M., Logan, J., Gladden, R. M., and Bohm, M. K. (2015). Vital signs: demographic and substance use trends among heroin users—united states, 2002–2013. MMWR. Morbidity and mortality weekly report 64, 719.

Karrer, B. and Newman, M. E. (2011). Stochastic blockmodels and community structure in networks. Physical review E 83, 016107.

McInnes, L., Healy, J., and Melville, J. (2018). Umap: Uniform manifold approximation and projection for dimension reduction. arXiv preprint 1802.03426.

McLachlan, G. J., Lee, S. X., and Rathnayake, S. I. (2019). Finite mixture models. Annual review of statistics and its application 6, 355–378.

McQuitty, L. L. (1966). Similarity analysis by reciprocal pairs for discrete and continuous data. Educational and Psychological measurement 26, 825–831.

Murtagh, F. and Legendre, P. (2014). Ward’s hierarchical agglomerative clustering method: which algorithms implement ward’s criterion? Journal of classification 31, 274–295.

NIH (2021). Overdose death rates. https://www.drugabuse.gov/drug-topics/trends-statistics/overdose-death-rates. Accessed: 2021-06-17.

Papastamoulis, P. (2016). label.switching: An R package for dealing with the label switching problem in MCMC outputs. Journal of Statistical Software 69, 1–24.

Peng, L. and Carvalho, L. (2016). Bayesian degree-corrected stochastic blockmodels for community detection. Electronic Journal of Statistics 10, 2746–2779.

Pons, P. and Latapy, M. (2005). Computing communities in large networks using random walks. In International symposium on computer and information sciences, pages 284–293. Springer.

R Core Team (2020). R: A Language and Environment for Statistical Computing. R Foundation for Statistical Computing, Vienna, Austria.

Richardson, N. R. and Roberts, D. C. (1996). Progressive ratio schedules in drug self-administration studies in rats: a method to evaluate reinforcing efficacy. Journal of neuroscience methods 66, 1–11.

Schwarz, G. et al. (1978). Estimating the dimension of a model. Annals of statistics 6, 461–464.

Shmulewitz, D., Greene, E. R., and Hasin, D. (2015). Commonalities and differences across substance use disorders: phenomenological and epidemiological aspects. Alcoholism: Clinical and Experimental Research 39, 1878–1900.

Snijders, T. A. and Nowicki, K. (1997). Estimation and prediction for stochastic blockmodels for graphs with latent block structure. Journal of classification 14, 75–100.

Stork, D. G., Duda, R. O., Hart, P. E., and Stork, D. (2001). Pattern classification. A Wiley-Interscience Publication.

Venniro, M., Banks, M. L., Heilig, M., Epstein, D. H., and Shaham, Y. (2020). Improving translation of animal models of addiction and relapse by reverse translation. Nature Reviews Neuroscience 21, 625–643.

Venniro, M., Zhang, M., Caprioli, D., Hoots, J. K., Golden, S. A., Heins, C., Morales, M., Epstein, D. H., and Shaham, Y. (2018). Volitional social interaction prevents drug addiction in rat models. Nature neuroscience 21, 1520–1529.

Woods, L. C. S. and Palmer, A. A. (2019). Using heterogeneous stocks for fine-mapping genetically complex traits. Rat Genomics pages 233–247.

